# Megadepth: efficient coverage quantification for BigWigs and BAMs

**DOI:** 10.1101/2020.12.17.423317

**Authors:** Christopher Wilks, Omar Ahmed, Daniel N. Baker, David Zhang, Leonardo Collado-Torres, Ben Langmead

## Abstract

**Motivation:** A common way to summarize sequencing datasets is to quantify data lying within genes or other genomic intervals. This can be slow and can require different tools for different input file types.

**Results:** Megadepth is a fast tool for quantifying alignments and coverage for BigWig and BAM/CRAM input files, using substantially less memory than the next-fastest competitor. Megadepth can summarize coverage within all disjoint intervals of the Gencode V35 gene annotation for more than 19,000 GTExV8 BigWig files in approximately one hour using 32 threads. Megadepth is available both as a command-line tool and as an R/Bioconductor package providing much faster quantification compared to the rtracklayer package.

**Availability:** https://github.com/ChristopherWilks/megadepth, https://bioconductor.org/packages/megadepth.

## 1 Introduction

Many sequencing data analyses are concerned with the depth of coverage in genomic regions. For example, RNA-seq alignments are often quantified within annotated intervals. Other examples include copy-number analysis of DNA-seq data or quantification of coverage under ChIP-seq peaks. The need is particularly pronounced for RNA-seq, where datasets may need periodic re-quantification with respect to updated or alternative gene annotations (Collado-Torres *et al.*, 2017).

BAM files store read alignments in a compressed and indexed form allowing random access (Li *et al.*, 2009). CRAM files are similar, additionally using reference-based compression (Hsi-Yang Fritz *et al.*, 2011). BigWig files (Kent *et al.*, 2010) store coverage vectors (not alignments) in a compressed and indexed form. While BAM and CRAM contain more information than BigWigs, BigWigs are also used for long-term storage because they are much smaller – often by an order of magnitude – while keeping enough information for requantification.

Mosdepth (Pedersen *et al.*, 2018) is an efficient quantification tool designed for BAM/CRAM files that can summarize coverage within intervals or across the entire file. Samtools and Sambamba (Li *et al.*, 2009; Tarasov *et al.*, 2015) can extract coverage from genomic regions within BAM and other related files (e.g. BED, VCF), though they cannot summarize coverage (e.g. sum or average). WiggleTools (Zerbino *et al.*, 2014) and bwtool (Pohl *et al.*, 2014) can extract and summarize coverage from BigWig files, and pyBigWig (Ramírez *et al.*, 2016) is a Python module with similar functionality. rtracklayer is an R/Biconductor package that handles both BAM and BigWig formats. In contrast, Megadepth supports BAM, CRAM, and BigWig inputs. It is faster while providing more features than other tools.

## 2 Methods

Megadepth processes BAMs one chromosome at a time, allocating a chromosome-length array in memory. It scans alignments in the BAM – possibly looking only within user-specified regions – and tallies base coverage in the array, either via the increment/decrement approach (Pedersen *et al.*, 2018; Wiewiórka *et al.*, 2019) or by storing explicit counts, depending on the operation. Megadepth uses the same general approach for BigWig files, scanning them base-by-base. Megadepth can output per-base coverage counts from BAM/CRAM inputs in a BED or BigWig file. Besides base-level coverage, Megadepth can additionally output per-interval coverage sums or averages as a BED file and an overall area-under-coverage (AUC) statistic. Megadepth can be configured to use multiple HTSlib threads for reading BAMs, speeding up block-gzip decompression (Supplementary Note 1). Since Megadepth’s single-threaded processing of BigWigs is already extremely fast (typical files take seconds) multi-threading is not implemented for that mode (Supplementary Note 2). Megadepth can query remote BAM, CRAM and BigWig files via an HTTP or FTP URL. Megadepth is written in C++11 and utilizes the HTSLib (v1.11) and libBigWig (v0.4.4) (Ramírez *et al.*, 2016) libraries. Binaries are available for Linux x86-64, MacOS x86-64, and Windows x86-64.

## 3 Results

We used BigWig-enabled tools to compute coverage sums for 5.5 million repetitive-element intervals across 10 BigWig files from GTEx brains (Table 1A). Megadepth was at least 4 times faster than all other tools while using 543 MiB of memory, the second lowest memory footprint among the 5 tools. WiggleTools was the next-fastest tool but it used *~*10 GiB of memory, limiting its utility on some systems. The megadepth-R package, which wraps Megadepth functionality for R, was 47 times faster and used a fraction of memory (808 MiB) compared to rtracklayer (*~*14 GiB), the only R/Bioconductor tool we tested. We performed more comparisons using different BigWigs files and intervals sets, including disjoint intervals from Gencode V35 (Supplementary Note 3). Overall, Megadepth was the fastest tool, though the speed gap was smaller for smaller interval sets; e.g. WiggleTools was only 30% slower for the Gencode V35 set. In addition, we recently used Megadepth to re-quantify all disjoint intervals of the Gencode V35 gene annotation for 19,214 GTExV8 BigWig files in about one hour using 32 threads.

**Table 1.** Top: Comparison of BigWig-enabled tools when computing coverage sums over repetitive-element intervals for 10 GTEx brain tissue BigWigs. Bottom: Comparison of BAM-enabled tools when computing coverage means over exome intervals for a 30X WGS BAM. Each tool’s features are also summarized.

| Tool | Relative Time | Run Time | Memory (MiB) | BAM Input | BigWig Input | MacOS | Windows Native | R Interface |
| --- | --- | --- | --- | --- | --- | --- | --- | --- |
| Megadepth (BigWig) | 1.00 | 1m:57s | 543 | yes | yes | yes | yes | yes |
| megadepth-R (BigWig) | 2.13 | 4m:09s | 808 | yes | yes | yes | yes | yes |
| WiggleTools | 4.06 | 7m:54s | 10,379 | no | yes | yes | no | no |
| pyBigWig | 68.13 | 2h:12m:36s | 7 | no | yes | yes | no | no |
| bwtool | 90.48 | 2h:56m:06s | 750 | no | yes | no | no | no |
| rtracklayer | 100.61 | 3h:15m:49s | 14,074 | yes | yes | yes | no | yes |
| Megadepth (BAM) | 1.00 | 2m:17s | 1,016 | yes | yes | yes | yes | yes |
| Mosdepth | 5.58 | 12m:43s | 1,911 | yes | no | yes | no | no |
| Samtools | 40.05 | 1h:31m:20s | 15 | yes | no | yes | yes | yes |
| Sambamba | 3.55 | 8m:05s | 157 | yes | no | yes | no | no |

Next we used the BAM-enabled tools to compute mean coverage within a set of 191,744 exome-capture intervals across a single 30X coverage whole-genome DNA-seq BAM (Table 1B). Megadepth was at least 3 times faster than other tools. While Megadepth used more memory (*~*1 GiB) compared to samtools and sambamba, it used about half the memory of the next-fastest tool, Mosdepth. Megadepth BAM processing is generally slower than BigWig processing since BAM files store substantially more information, e.g. including read sequences and base qualities. Supplementary Note 4 describes comparisons on BAM and CRAM files where the tools are configured to output base-by-base coverage values. While Megadepth is still fastest, some of the differences are very small, e.g. Mosdepth is only 12% slower. But the difference grows when using a RNA-seq BAM file, where Mosdepth takes 2.7x the time. We also measured the time required to analyze an entire DNA-seq BAM file within 500 bp windows, similar to a benchmark in the Mosdepth study (Supplementary Note 5). Finally, we performed further BAM and CRAM benchmarks using query intervals (Supplementary Note 6).

## 4 Discussion

Megadepth is an efficient tool for quantifying alignments and coverage within genomic intervals. It handles BigWig, BAM and CRAM files at faster speeds than any other tool, and with lower memory footprint than the next-fastest tools. Quantification is a common way to analyze new datasets and to re-analyze archived sequencing datasets (Zhang *et al.*, 2020; Collado-Torres *et al.*, 2017). Megadepth further facilities this by providing an R/Bioconductor interface, readily used in combination with recount2 and other R-based resources. BigWig support is of particular import since BigWigs are much smaller than BAMs, while still containing the information needed to re-quantify. Megadepth facilitates this both by enabling rapid conversion from BAM to BigWig – a onetime cost – and by rapidly re-quantifying the resulting BigWig with respect to newer interval sets, possibly many times. Finally, Megadepth supports extraction of alternate base coverage, junction co-occurrences, and fragment length distribution for paired samples (Supplementary Figure S3).

## Funding

CW, OA, DNB and BL were supported by NIH/NIGMS grant R01GM118568 to BL. LCT, BL and CW were supported by R01GM121459 to Dr. Kasper Hansen. DZ was supported by UK Medical Research Council funding awarded to Dr. Mina Ryten (Tenure Track Clinician Scientist Fellowship, MR/N008324/1).

## Notes

https://github.com/ChristopherWilks/megadepth

https://bioconductor.org/packages/megadepth

### Competing Interest Statement

The authors have declared no competing interest.

