## Supplemental Data for "Megadepth: efficient coverage quantification for BigWigs and BAMs"

*Version Note*

The versions of the software used throughout the benchmarks in this set of supplementary notes are:

- Megadepth 1.0.9 (using HTSLib 1.11 and libBigWig 0.4.4)
- Mosdepth 0.3.1 (statically compiled by these authors to use HTSLib 1.11)
- Sambamba 0.7.1
- Samtools 1.11
- WiggleTools 1.2.4
- pyBigWig 0.3.17
- bwtool 1.0
- rtracklayer 1.50.0

All benchmarks were run on a machine with four Intel Xeon E7-4830 v4 (Broadwell) 2.00 GHz processors (112 threads in total) and 1 TB of RAM. Experiments read from/wrote to a direct-attached storage array.

*Windows Native Version Note*

In Table 1 of the main manuscript, the term "Windows Native" refers to a pre-compiled binary that does not require Windows Subsystem for Linux (WSL), Cygwin, or similar compatibility layer. Megadepth does not support writing BigWig files on Windows at this time.

---

*Supplementary Note S1: Comparing megadepth and mosdepth's scalability with additional BAM decompression threads*

**Description:** This experiment looks at time savings that occur when trying to compute mean coverage across regions on BAMs with each additional thread. This experiment does it for both an annotation consisting of 500 bp sliding windows and hg19 exome annotation.

**Data:**

- Genomic BAM:
  - Used ERR1019041.bam from Simons Genome Diversity Panel (Mallick et al., 2016)
  - Data was preprocessed with following commands:
    - samtools view -b -s 12.75 ERR1019041.bam > ERR1019041_sub.bam
    - samtools sort ERR1019041_sub.bam > ERR1019041_sorted.bam
    - samtools -b view chr1 … chr22 chrX chrY > ERR1019041_sorted_minusM.bam
    - samtools index ERR1019041_sorted.bam
  - For reference, this BAM is 69 GB
- Used hg19 exome from this link: https://www.twistbioscience.com/resources/bed-file/ngs-human-core-exome-panel-bed-files

**Sub-Experiments:**

*Table S1a: Looking at time improvements with each additional BAM decompression thread when computing mean coverage of Genomic BAM file (ERR1019041) with sliding window regions of 500 bp*

| Format | Tool | Additional Threads | Command |
|---|---|---|---|
| BAM | megadepth | t = 0:8 | megadepth <bam_file.bam> --threads <t> --gzip --op mean --annotation 500 --prefix <output_prefix> --no-annotation-stdout |
| BAM | mosdepth | t = 0:8 | mosdepth -F 260 -n --by 500 --threads < t> <output_prefix> <bam_file.bam> |

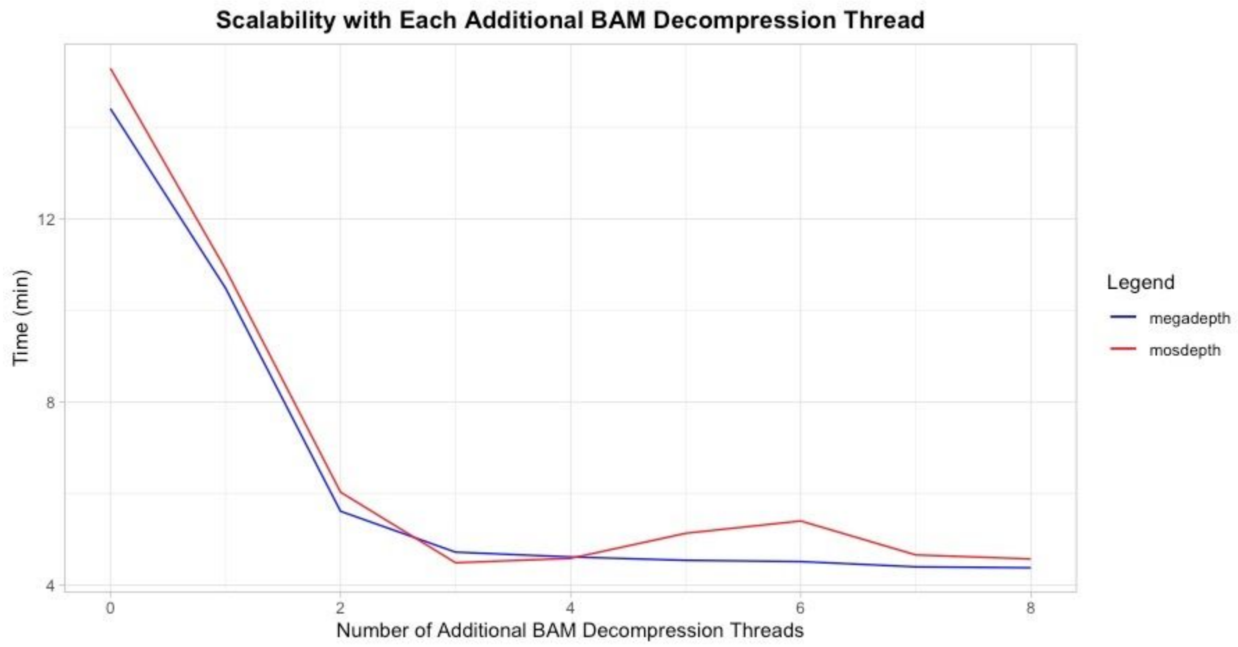


Figure S1. Thread scaling of whole-genome base coverage across a BAM

*Table S1b: Looking at time improvements with each additional BAM decompression thread on computing mean on regions for Genomic BAM file (ERR1019041) with hg19 exome annotation*

| Format | Tool | Additional Threads | Command |
|---|---|---|---|
| BAM | megadepth | t = 0:8 | megadepth <bam_file.bam> --threads < t> --gzip --op mean --annotation <bed_file.bed> --prefix <output_prefix> --no-annotation-stdout |
| BAM | mosdepth | t = 0:8 | mosdepth -F 260 -n --by <bed_file.bed> --threads <t> <output_prefix> <bam_file.bam> |

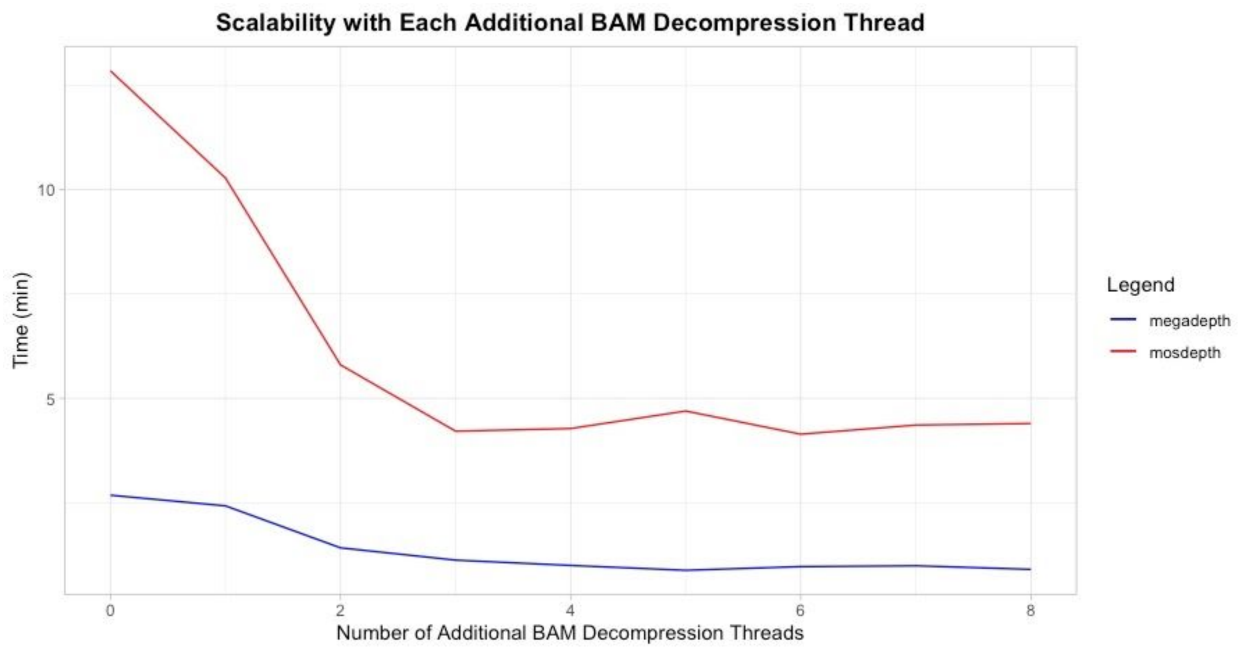


Figure S2. Thread scaling of exome base coverage across a BAM

---

*Supplementary Note S2: Looking at megadepth's ability to process all the BigWigs produced from GTEx*

**Description:** Examine the time and memory needed for megadepth to compute the sums on regions for all the BigWig files produced from GTEx*.  We used the GNU parallel (Tange 2011) command to run multiple instances of Megadepth concurrently on the same system.

**Data:**

- Input data used was BigWig files produced from GTEx (19,214 BigWig files), available through recount3 project
- Annotation used was Gencodev35 (Frankish et al., 2019) disjoint exon

**Sub-Experiments:**

*Table S2a: Examining time and memory used by megadepth to compute sum of regions on all GTEx BigWigs on GencodeV35 disjoint exon regions*

| Format | Tool | Command | Jobs | Total Run Time |
|---|---|---|---|---|
| BigWig | megadepth | bash command_list.txt | 1 | 21:26:24 |
| BigWig | megadepth | parallel -j 4 < command_list.txt | 4 | 05:14:28 |
| BigWig | megadepth | parallel -j 8 < command_list.txt | 8 | 02:45:25 |

| | | | | |
|---|---|---|---|---|
| BigWig | megadepth | parallel -j 16 < command_list.txt | 16 | 01:30:39 |
| BigWig | megadepth | parallel -j 32 < command_list.txt | 32 | 01:05:15 |

*The Genotype-Tissue Expression (GTEx) Project was supported by the Common Fund of the Office of the Director of the National Institutes of Health, and by NCI, NHGRI, NHLBI, NIDA, NIMH, and NINDS. The data used for the analyses described in this manuscript were obtained from: the GTEx Portal on 11/2019 and dbGaP accession number phs000424.v8.p2 on 11/2019.

---

*Supplementary Note S3: Comparing ability to compute sums on regions of BigWigs*

**Description:** Compare the ability of megadepth to compute the sum of coverage on regions of BigWig files.

**Data:**

- Input data used was BigWig files produced from GTEx, available through recount project
- Annotations used were Gencodev35 disjoint exon and repetitive element annotation

**Sub-Experiments:**

*Table S3a: Comparing ability to compute sum of coverage on 5 BRAIN GTEx files across annotated regions in Gencodev35 disjoint exons*

| Format | Tool | Command | Relative Time | Total Run Time | Average Memory (MiB) |
|---|---|---|---|---|---|
| BigWig | megadepth | megadepth <bigwig_file.bw> --annotation <bed_file.bed> --op sum > <output_file> | 1.000 | 00:00:18 | 106.09 |
| BigWig | wiggletools | wiggletools apply_paste <output_file> AUC <bed_file.bed> <bigwig_file.bw> | 1.303 | 00:00:24 | 762.54 |
| BigWig | megadepth-R | megadepth::get_coverage(bigwig_file=bw, annotation=annotation_path, op = “sum”) | 2.778 | 00:00:50 | 426 |
| BigWig | pyBigWig | python pybig_benchmark.py <bed_file.bed> <bigwig_file.bw> <output_file> | 26.891 | 00:08:09 | 7.28 |
| BigWig | bwtool | bwtool summary <bed_file.bed> <bigwigfile.bw> -with-sum -keep-bed -fill=0 -header <output_file> | 35.871 | 00:10:52 | 95.78 |
| BigWig | rtracklayer | Rscript get_coverage.R <bed_file.bed> <big_wig.bw> | 45.081 | 00:13:40 | 2858.31 |

*Table S3b: Comparing ability to compute sum of coverage on 5 BRAIN GTEx files across annotated regions in repetitive elements*

| Format | Tool | Command | Relative Time | Total Run Time | Average Memory (MiB) |
|---|---|---|---|---|---|
| BigWig | megadepth | megadepth <bigwig_file.bw> --annotation <bed_file.bed> --op sum > <output_file> | 1.000 | 00:00:49 | 541.12 |
| BigWig | wiggletools | wiggletools apply_paste <output_file> AUC <bed_file.bed> <bigwig_file.bw> | 4.098 | 00:03:21 | 10379.14 |
| BigWig | megadepth-R | megadepth::get_coverage(bigwig_file=bw, annotation=annotation_path, op = “sum”) | 2.612 | 00:02:08 | 808 |
| BigWig | pyBigWig | python pybig_benchmark.py <bed_file.bed> <bigwig_file.bw> <output_file> | 78.930 | 01:04:23 | 7.24 |
| BigWig | bwtool | bwtool summary <bed_file.bed> <bigwigfile.bw> -with-sum -keep-bed -fill=0 -header <output_file> | 103.722 | 01:24:36 | 749.68 |
| BigWig | rtracklayer | Rscript get_coverage.R <bed_file.bed> <big_wig.bw> | 105.889 | 01:26:22 | 14074.47 |

*Table S3c: Comparing ability to compute sum of coverage on 10 BRAIN GTEx files across annotated regions in Gencodev35 disjoint exons*

| Format | Tool | Command | Relative Time | Total Run Time | Average Memory (MiB) |
|---|---|---|---|---|---|
| BigWig | megadepth | megadepth <bigwig_file.bw> --annotation <bed_file.bed> --op sum > <output_file> | 1.000 | 00:00:36 | 101 |
| BigWig | wiggletools | wiggletools apply_paste <output_file> AUC <bed_file.bed> <bigwig_file.bw> | 1.577 | 00:00:57 | 757 |
| BigWig | megadepth-R | megadepth::get_coverage(bigwig_file=bw, annotation=annotation_path, op = “sum”) | 2.778 | 00:01:40 | 426 |
| BigWig | pyBigWig | python pybig_benchmark.py <bed_file.bed> <bigwig_file.bw> <output_file> | 29.519 | 00:17:45 | 7 |
| BigWig | bwtool | bwtool summary <bed_file.bed> <bigwigfile.bw> -with-sum -keep-bed -fill=0 -header <output_file> | 38.906 | 00:23:24 | 96 |
| BigWig | rtracklayer | Rscript get_coverage.R <bed_file.bed> <big_wig.bw> | 46.962 | 00:28:15 | 2824 |

*Table S3d: Comparing ability to compute sum of coverage on 10 BRAIN GTEx files across annotated regions in repetitive elements*

| Format | Tool | Command | Relative Time | Total Run Time | Average Memory (MiB) |
|---|---|---|---|---|---|
| BigWig | megadepth | megadepth <bigwig_file.bw> --annotation <bed_file.bed> --op sum > <output_file> | 1.000 | 00:01:57 | 543 |
| BigWig | wiggletools | wiggletools apply_paste <output_file> AUC <bed_file.bed> <bigwig_file.bw> | 4.062 | 00:07:54 | 10379 |
| BigWig | megadepth-R | megadepth::get_coverage(bigwig_file=bw, annotation=annotation_path, op = "sum") | 2.128 | 00:04:09 | 808 |
| BigWig | pyBigWig | python pybig_benchmark.py <bed_file.bed> <bigwig_file.bw> <output_file> | 68.127 | 02:12:36 | 7 |
| BigWig | bwtool | bwtool summary <bed_file.bed> <bigwigfile.bw> -with-sum -keep-bed -fill=0 -header <output_file> | 90.480 | 02:56:06 | 750 |
| BigWig | rtracklayer | Rscript get_coverage.R <bed_file.bed> <big_wig.bw> | 100.611 | 03:15:49 | 14074 |

*Table S3e: Summary statistics of the two annotations used in Section 5*

| Annotation File | Number of Regions | Min | Max | Mean | Median |
|---|---|---|---|---|---|
| Gencodev35 Disjoint Exons | 665,224 | 1 | 320,039 | 227 | 98 |
| Repetitive Elements | 5,520,278 | 19 | 500,000 | 286 | 190 |

---

<u>*Supplementary Note S4: Comparing ability to output coverage per-base coverage on BAMs/CRAMs*</u>

**Description:** Compare the ability of megadepth to output the per-base coverage of BAM/CRAM files with other tools that have that same functionality. This mode will produce coverage across a whole genome or transcriptome depending on the input.

**Data:**

- Genomic BAM from Supplementary Note S1 (ERR1019041)
- RNA-seq BAM:

    - Geuvadis RNA-seq BAM (Lappalainen 2013), ERR204876_ERP001942_hg38.sorted.bam
    - For reference, this BAM is 716 MiB
- CRAM File from 1KGP (Auton et al., 2015):
    - ERR3239958
    - For reference, this CRAM is 14 GB

**Sub-Experiments:**

*Table S4a: Comparing ability to compute per-base coverage on Genomic BAM file with single thread (ERR1019041)*

| Format | Tool | Command | Relative Time | Run Time | Memory (MiB) |
|---|---|---|---|---|---|
| BAM | megadepth | megadepth <bam_file.bam> --gzip --no-coverage-stdout --coverage --prefix <ouput_prefix> | 1.000 | 00:16:59 | 949 |
| BAM | mosdepth | mosdepth -F 260 <output_prefix> <bam_file.bam> | 1.156 | 00:19:38 | 1904 |
| BAM | samtools | samtools depth -a -G 260 -o <output_file_name> <bam_file.bam> | 3.333 | 00:56:37 | 10 |
| BAM | sambamba | sambamba depth base -o <output_file> -c 0 <bam_file.bam> | 6.518 | 01:50:44 | 221 |
| BAM | bedtools | bedtools genomecov -d -split -ibam <bam_file.bam> > <output_file> | 7.484 | 02:07:09 | 1858 |

*Table S4b: Comparing ability to compute per-base coverage on RNA-seq BAM file with single thread (ERR204876_ERP001942)*

| Format | Tool | Command | Relative Time | Run Time | Memory (MiB) |
|---|---|---|---|---|---|
| BAM | megadepth | megadepth <bam_file.bam> --gzip --no-coverage-stdout --coverage --prefix <ouput_prefix> | 1.000 | 00:00:24 | 940 |
| BAM | mosdepth | mosdepth -F 260 <output_prefix> <bam_file.bam> | 2.687 | 00:01:05 | 1871 |
| BAM | samtools | samtools depth -a -G 260 -o <output_file_name> <bam_file.bam> | 41.953 | 00:16:52 | 7 |
| BAM | sambamba | sambamba depth base -o <output_file> -c 0 <bam_file.bam> | 78.216 | 00:31:27 | 51 |
| BAM | bedtools | bedtools genomecov -d -split -ibam <bam_file.bam> > <output_file> | 244.548 | 01:38:19 | 1856 |

*Table S4c: Comparing ability to compute per-base coverage on Genomic CRAM file with single thread (ERR3239958)*

| Format | Tool | Command | Relative Time | Run Time | Memory (MiB) |
|---|---|---|---|---|---|
| CRAM | megadepth | megadepth <cram_file.cram> --gzip --fasta <fasta_ref> --no-coverage-stdout --coverage --prefix <ouput_prefix> | 1.000 | 00:11:03 | 986 |
| CRAM | mosdepth | mosdepth -F 260 -f <fasta_ref> <output_prefix> <cram_file.cram> | 1.275 | 00:14:06 | 1912 |
| CRAM | samtools | samtools depth -a -G 260 -o <output_file> <cram_file.cram> | 4.271 | 00:47:13 | 87 |

---

### *Supplementary Note S5: Comparing ability to output mean coverage on a sliding window over the genome*

**Description:** Compare the ability of megadepth to output the average coverage on sliding windows of BAM files with other tools that have that same functionality.

**Data:**
- Used the Genomic BAM from Supplementary Note S1 (ERR1019041)

**Sub-Experiments:**

*Table S5a: Comparing ability to compute mean coverage on Genomic BAM file (ERR1019041) in 500 bp sliding windows with single thread (run time is mean of 5 runs)*

| Format | Tool | Command | Relative Time | Average Run Time | Memory (MiB) |
|---|---|---|---|---|---|
| BAM | megadepth | megadepth <bam_file.bam> --gzip --op mean --annotation 500 --prefix <output_prefix> --no-annotation-stdout | 1.000 | 00:11:33 | 950 |
| BAM | mosdepth | mosdepth -F 260 -n --by 500 <output_prefix> <bam_file.bam> | 1.115 | 00:12:53 | 1906 |
| BAM | samtools | samtools -G 260 -b <bed_file> -o <output_file> <bam_file.bam> | 4.973 | 00:57:26 | 105 |
| BAM | sambamba | sambamba depth region --regions=<bed_file.bed> -o <output_file> <bam_file.bam> | 5.251 | 01:00:39 | 1566 |

---

*Supplementary Note S6: Comparing ability to output mean coverage on regions of BAMs/CRAMs*

**Description:** Compare the ability of megadepth to output the average coverage on given intervals of BAM and CRAM files with other tools that have that same functionality.

**Data:**

- Input Data:
    - Used Genomic BAM (ERR1019041) from Supplementary Note S1 as well as Genomic CRAM (ERR3239958) and RNA-seq BAM (ERR204876_ERP001942) from Supplementary Note S4
- Annotations:
    - Took Gencodev35 GTF annotation file used rtracklayer to help convert it into a GFF file with disjoint exons, then used a BEDOPs script to convert to bed file that will be used with programs.
    - Used annotation for repetitive elements.
    - Used hg19 exome annotation from Supplementary Note S1 for Simons Genomic BAM in Experiment 6a and 6b
    - Used hg38 exome annotation for 1KGP CRAM in Experiments S6g and S6h
        - Both exome annotations obtained at this link: https://www.twistbioscience.com/resources/bed-file/ngs-human-core-exome-panel-bed-files

**Sub-Experiments:**

*Table S6a: Comparing ability to compute mean coverage on Genomic BAM file (ERR1019041) in hg19 exome regions (1 thread)*

| Format | Tool | Threads | Command | Relative Time | Run Time | Memory (MiB) |
|---|---|---|---|---|---|---|
| BAM | megadepth | 1 | megadepth <bam_file.bam> --gzip --op mean --annotation <bed_file.bed> --prefix <output_prefix> --no-annotation-stdout | 1.000 | 00:02:17 | 1016 |
| BAM | mosdepth | 1 | mosdepth -F 260 -n --by <bed_file.bed> <output_prefix> <bam_file.bam> | 5.579 | 00:12:43 | 1911 |
| BAM | samtools | 1 | samtools -G 260 -b <bed_file.bed> -o <output_file> <bam_file.bam> | 40.051 | 01:31:20 | 15 |
| BAM | sambamba | 1 | sambamba depth region --regions=<bed_file.bed> -o <output_file> <bam_file.bam> | 3.547 | 00:08:05 | 157 |

*Table S6b: Comparing ability to compute mean coverage on Genomic BAM file (ERR1019041) in hg19 exome regions (4 threads)*

| Format | Tool | Threads | Command | Relative Time | Run Time | Memory (MiB) |
|---|---|---|---|---|---|---|
| BAM | megadepth | 4 | megadepth <bam_file.bam> --threads 3 --gzip --op mean --annotation <bed_file.bed> --prefix <output_prefix> --no-annotation-stdout | 1.000 | 00:01:05 | 1017 |
| BAM | mosdepth | 4 | mosdepth -F 260 -n --threads 3 --by <bed_file.bed> --threads 3 <output_prefix> <bam_file.bam> | 3.907 | 00:04:14 | 1913 |
| BAM | sambamba | 4 | sambamba depth region --regions=<bed_file.bed> -t 4 -o <output_file> <bam_file.bam> | 2.294 | 00:02:29 | 134 |

*Table S6c: Comparing ability to compute mean coverage on RNA-seq BAM file (ERR204876_ERP001942) in Gencodev35 disjoint exon regions (1 thread)*

| Format | Tool | Threads | Command | Relative Time | Run Time | Memory (MiB) |
|---|---|---|---|---|---|---|
| BAM | megadepth | 1 | megadepth <bam_file.bam> --gzip --op mean --annotation <bed_file.bed> --prefix <output_prefix> --no-annotation-stdout | 1.000 | 00:00:23 | 1054 |
| BAM | mosdepth | 1 | mosdepth -F 260 -n --by <bed_file.bed> <output_prefix> <bam_file.bam> | 2.335 | 00:00:53 | 1974 |
| BAM | samtools | 1 | samtools -G 260 -b <bed_file> -o <output_file> <bam_file> | 15.037 | 00:05:42 | 19 |
| BAM | sambamba | 1 | sambamba depth region --regions=<bed_file.bed> -o <output_file> <bam_file.bam> | 17.731 | 00:06:43 | 1011 |

*Table S6d: Comparing ability to compute mean coverage on RNA-seq BAM file (ERR204876_ERP001942) in Gencodev35 disjoint exon regions (4 threads)*

| Format | Tool | Threads | Command | Relative Time | Run Time | Memory (MiB) |
|---|---|---|---|---|---|---|
| BAM | megadepth | 4 | megadepth <bam_file.bam> --threads 3 --gzip --op mean --annotation <bed_file.bed> --prefix <output_prefix> --no-annotation-stdout | 1.000 | 00:00:19 | 1055 |
| BAM | mosdepth | 4 | mosdepth -F 260 -n --threads 3 --by <bed_file.bed> <output_prefix> <bam_file.bam> | 2.251 | 00:00:44 | 1975 |
| BAM | sambamba | 4 | sambamba depth region --regions=<bed_file.bed> -t 4 -o <output_file> <bam_file.bam> | 21.211 | 00:06:52 | 1000 |

*Table S6e: Comparing ability to compute mean coverage on RNA-seq BAM file (ERR204876_ERP001942) in repetitive element regions (1 thread)*

| Format | Tool | Threads | Command | Relative Time | Run Time | Memory (MiB) |
|---|---|---|---|---|---|---|
| BAM | megadepth | 1 | megadepth <bam_file.bam> --gzip --op mean --annotation <bed_file> --prefix <output_prefix> --no-annotation-stdout | 1.000 | 00:00:39 | 1951 |
| BAM | mosdepth | 1 | mosdepth -F 260 -n --by <bed_file.bed> <output_prefix> <bam_file.bam> | 1.975 | 00:01:17 | 2442 |
| BAM | samtools | 1 | samtools -G 260 -b <bed_file> -o <output_file> <bam_file> | 12.639 | 00:08:15 | 92 |
| BAM | sambamba | 1 | sambamba depth region --regions=<bed_file.bed> -o <output_file> <bam_file.bam> | 11.230 | 00:07:20 | 1875 |

*Table S6f: Comparing ability to compute mean coverage on RNA-seq BAM file (ERR204876_ERP001942) in repetitive elements regions (4 threads)*

| Format | Tool | Threads | Command | Relative Time | Run Time | Memory (MiB) |
|---|---|---|---|---|---|---|
| BAM | megadepth | 4 | megadepth <bam_file.bam> --threads 3 --gzip --op mean --annotation <bed_file.bed> --prefix <output_prefix> --no-annotation-stdout | 1.000 | 00:00:35 | 1952 |
| BAM | mosdepth | 4 | mosdepth -F 260 -n --threads 3 --by <bed_file.bed> <output_prefix> <bam_file.bam> | 2.262 | 00:01:19 | 2446 |
| BAM | sambamba | 4 | sambamba depth region --regions=<bed_file.bed> -t 4 -o <output_file> <bam_file.bam> | 12.374 | 00:07:13 | 1842 |

*Table S6g: Comparing ability to compute mean coverage on Genomic CRAM (ERR3239958) in hg38 exome regions (1 thread)*

| Format | Tool | Threads | Command | Relative Time | Run Time | Memory (MiB) |
|---|---|---|---|---|---|---|
| BAM | megadepth | 1 | megadepth <cram_file.cram> --fasta <fasta_ref> --gzip --op mean --annotation <bed_file.bed> --prefix <output_prefix> --no-annotation-stdout | 1.000 | 00:02:16 | 1045 |
| BAM | mosdepth | 1 | mosdepth -F 260 -n -f <fasta_ref> --by <bed_file> <output_prefix> <cram_file.bam> | 3.464 | 00:07:50 | 1926 |
| BAM | samtools | 1 | samtools -G 260 -b <bed_file> -o <output_file> <cram_file> | 38.076 | 01:26:05 | 92 |

*Table S6h: Comparing ability to compute mean coverage on Genomic CRAM (ERR3239958) in hg38 exome regions (4 threads)*

| Format | Tool | Threads | Command | Relative Time | Run Time | Memory (MiB) |
|---|---|---|---|---|---|---|
| BAM | megadepth | 4 | megadepth <cram_file.cram> --fasta <fasta_ref> --threads 3 --gzip --op mean --annotation <bed_file.bed> --prefix <output_prefix> --no-annotation-stdout | 1.000 | 00:01:32 | 1049 |
| BAM | mosdepth | 4 | mosdepth -F 260 -n --threads 3 -f <fasta_ref> --by <bed_file> <output_prefix> <cram_file.bam> | 2.806 | 00:04:17 | 1993 |

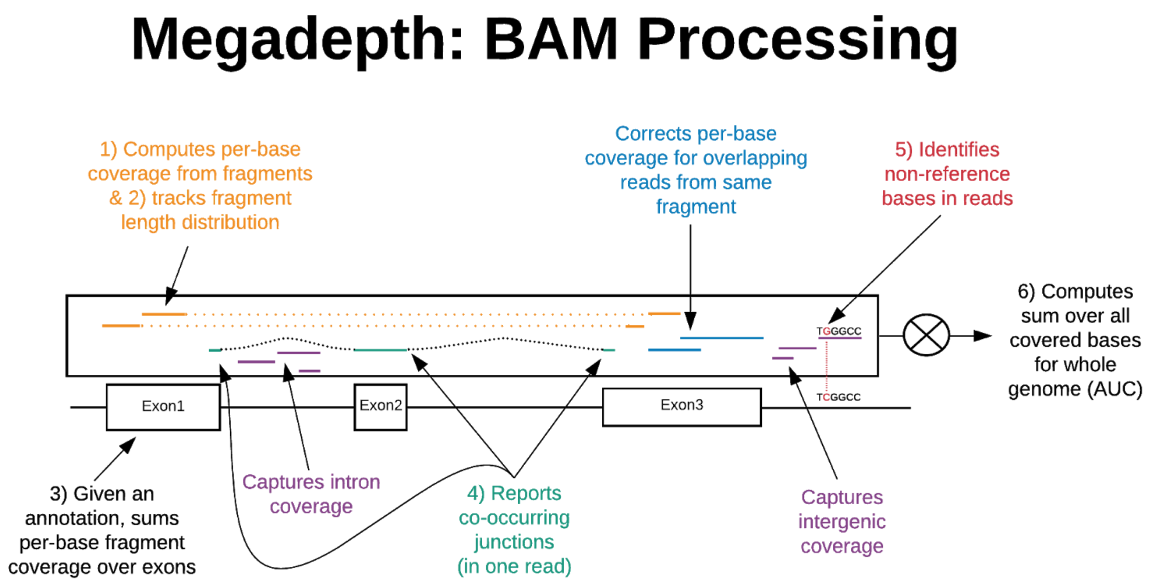


Figure S3. Coverage-related operations Megadepth can perform over a BAM file

Auton, A, Brooks, LD, Durbin, RM, Garrison, EP, Kang, HM, Korbel, JO, Marchini, JL, McCarthy, S, McVean, GA, Abecasis, GR, Auton, A, Abecasis, GR, Altshuler, DM, Durbin, RM, Abecasis, GR, Bentley, DR, Chakravarti, A, Clark, AG, Donnelly, P, Eichler, EE, Flicek, P, Gabriel, SB, Gibbs, RA, Green, ED, Hurles, ME, Knoppers, BM, Korbel, JO, Lander, ES, Lee, C, Lehrach, H, Mardis, ER, Marth, GT, McVean, GA, Nickerson, DA, Schmidt, JP, Sherry, ST, Wang, J, Wilson, RK, Gibbs, RA, Boerwinkle, E, Doddapaneni, H, Han, Y, Korchina, V, Kovar, C, Lee, S, Muzny, D, Reid, JG, Zhu, Y, Wang, J, Chang, Y, Feng, Q, Fang, X, Guo, X, Jian, M, Jiang, H, Jin, X, Lan, T, Li, G, Li, J, Li, Y, Liu, S, Liu, X, Lu, Y, Ma, X, Tang, M, Wang, B, Wang, G, Wu, H, Wu, R, Xu, X, Yin, Y, Zhang, D, Zhang, W, Zhao, J, Zhao, M, Zheng, X, Lander, ES, Altshuler, DM, Gabriel, SB, Gupta, N, Gharani, N, Toji, LH, Gerry, NP, Resch, AM, Flicek, P, Barker, J, Clarke, L, Gil, L, Hunt, SE, Kelman, G, Kulesha, E, Leinonen, R, McLaren, WM,

Radhakrishnan, R, Roa, A, Smirnov, D, Smith, RE, Streeter, I, Thormann, A, Toneva, I, Vaughan, B, Zheng-Bradley, X, Bentley, DR, Grocock, R, Humphray, S, James, T, Kingsbury, Z, Lehrach, H, Sudbrak, R, Albrecht, MW, Amstislavskiy, VS, Borodina, TA, Lienhard, M, Mertes, F, Sultan, M, Timmermann, B, Yaspo, ML, Mardis, ER, Wilson, RK, Fulton, L, Fulton, R, Sherry, ST, Ananiev, V, Belaia, Z, Beloslyudtsev, D, Bouk, N, Chen, C, Church, D, Cohen, R, Cook, C, Garner, J, Hefferon, T, Kimelman, M, Liu, C, Lopez, J, Meric, P, O'Sullivan, C, Ostapchuk, Y, Phan, L, Ponomarov, S, Schneider, V, Shekhtman, E, Sirotkin, K, Slotta, D, Zhang, H, McVean, GA, Durbin, RM, Balasubramaniam, S, Burton, J, Danecek, P, Keane, TM, Kolb-Kokocinski, A, McCarthy, S, Stalker, J, Quail, M, Durbin, RM, Balasubramaniam, S, Burton, J, Danecek, P, Keane, TM, Kolb-Kokocinski, A, McCarthy, S, Stalker, J, Quail, M, Schmidt, JP, Davies, CJ, Gollub, J, Webster, T, Wong, B, Zhan, Y, Auton, A, Campbell, CL, Kong, Y, Marcketta, A, Gibbs, RA, Yu, F, Antunes, L, Bainbridge, M, Muzny, D, Sabo, A, Huang, Z, Wang, J, Coin, LJ, Fang, L, Guo, X, Jin, X, Li, G, Li, Q, Li, Y, Li, Z, Lin, H, Liu, B, Luo, R, Shao, H, Xie, Y, Ye, C, Yu, C, Zhang, F, Zheng, H, Zhu, H, Alkan, C, Dal, E, Kahveci, F, Marth, GT, Garrison, EP, Kural, D, Lee, WP, Leong, WF, Stromberg, M, Ward, AN, Wu, J, Zhang, M, Daly, MJ, DePristo, MA, Handsaker, RE, Altshuler, DM, Banks, E, Bhatia, G, Del Angel, G, Gabriel, SB, Genovese, G, Gupta, N, Li, H, Kashin, S, Lander, ES, McCarroll, SA, Nemesh, JC, Poplin, RE, Yoon, SC, Lihm, J, Makarov, V, Clark, AG, Gottipati, S, Keinan, A, Rodriguez-Flores, JL, Korbel, JO, Rausch, T, Fritz, MH, StÃ¼tz, AM, Flicek, P, Beal, K, Clarke, L, Datta, A, Herrero, J, McLaren, WM, Ritchie, GR, Smith, RE, Zerbino, D, Zheng-Bradley, X, Sabeti, PC, Shlyakhter, I, Schaffner, SF, Vitti, J, Cooper, DN, Ball, EV, Stenson, PD, Bentley, DR, Barnes, B, Bauer, M, Cheetham, RK, Cox, A, Eberle, M, Humphray, S, Kahn, S, Murray, L, Peden, J, Shaw, R, Kenny, EE, Batzer, MA, Konkel, MK, Walker, JA, MacArthur, DG, Lek, M, Sudbrak, R, Amstislavskiy, VS, Herwig, R, Mardis, ER, Ding, L, Koboldt, DC, Larson, D, Ye, K, Gravel, S, Swaroop, A, Chew, E, Lappalainen, T, Erlich, Y, Gymrek, M, Willems, TF, Simpson, JT, Shriver, MD, Rosenfeld, JA, Bustamante, CD, Montgomery, SB, De La Vega, FM, Byrnes, JK, Carroll, AW, DeGorter, MK, Lacroute, P, Maples, BK, Martin, AR, Moreno-Estrada, A, Shringarpure, SS, Zakharia, F, Halperin, E, Baran, Y, Lee, C, Cerveira, E, Hwang, J, Malhotra, A, Plewczynski, D, Radew, K, Romanovitch, M, Zhang, C, Hyland, FC, Craig, DW, Christoforides, A, Homer, N, Izatt, T, Kurdoglu, AA, Sinari, SA, Squire, K, Sherry, ST, Xiao, C, Sebat, J, Antaki, D, Gujral, M, Noor, A, Ye, K, Burchard, EG, Hernandez, RD, Gignoux, CR, Haussler, D, Katzman, SJ, Kent, WJ, Howie, B, Ruiz-Linares, A, Dermitzakis, ET, Devine, SE, Abecasis, GR, Kang, HM, Kidd, JM, Blackwell, T, Caron, S, Chen, W, Emery, S, Fritsche, L, Fuchsberger, C, Jun, G, Li, B, Lyons, R, Scheller, C, Sidore, C, Song, S, Sliwerska, E, Taliun, D, Tan, A, Welch, R, Wing, MK, Zhan, X, Awadalla, P, Hodgkinson, A, Li, Y, Shi, X, Quitadamo, A, Lunter, G, McVean, GA, Marchini, JL, Myers, S, Churchhouse, C, Delaneau, O, Gupta-Hinch, A, Kretzschmar, W, Iqbal, Z, Mathieson, I, Menelaou, A, Rimmer, A, Xifara, DK, Oleksyk, TK, Fu, Y, Liu, X, Xiong, M, Jorde, L, Witherspoon, D, Xing, J, Eichler, EE, Browning, BL, Browning, SR, Hormozdiari, F, Sudmant, PH, Khurana, E, Durbin, RM, Hurles, ME, Tyler-Smith, C, Albers, CA, Ayub, Q, Balasubramaniam, S, Chen, Y, Colonna, V, Danecek, P, Jostins, L, Keane, TM, McCarthy, S, Walter, K, Xue, Y, Gerstein, MB, Abyzov, A, Balasubramanian, S, Chen, J, Clarke, D, Fu, Y, Harmanci, AO, Jin, M, Lee, D, Liu, J, Mu, XJ, Zhang, J, Zhang, Y, Gerstein, MB, Abyzov, A, Balasubramanian, S, Chen, J, Clarke, D, Fu, Y, Harmanci, AO, Jin, M, Lee, D, Liu, J, Mu, XJ, Zhang, J, Zhang, Y, Li, Y, Luo, R, Zhu, H, Alkan, C, Dal, E, Kahveci, F, Marth, GT, Garrison, EP, Kural, D, Lee, WP, Ward, AN, Wu, J, Zhang, M, McCarroll, SA, Handsaker, RE, Altshuler, DM, Banks, E, Del Angel, G, Genovese, G, Hartl, C, Li, H, Kashin, S, Nemesh, JC, Shakir, K, Yoon, SC, Lihm, J, Makarov, V, Degenhardt, J, Korbel, JO, Fritz, MH, Meiers, S, Raeder, B, Rausch, T, StÃ¼tz, AM, Flicek, P, Casale, FP, Clarke, L, Smith, RE, Stegle, O, Zheng-Bradley, X, Bentley, DR, Barnes, B, Cheetham, RK, Eberle, M, Humphray, S, Kahn, S, Murray, L, Shaw, R, Lameijer, EW, Batzer, MA, Konkel, MK, Walker, JA, Ding, L, Hall, I, Ye, K, Lacroute, P, Lee, C, Cerveira, E, Malhotra, A, Hwang, J, Plewczynski, D, Radew, K, Romanovitch, M, Zhang, C, Craig, DW, Homer, N, Church, D, Xiao, C, Sebat, J, Antaki, D, Bafna, V, Michaelson, J, Ye, K, Devine, SE, Gardner, EJ, Abecasis, GR, Kidd, JM, Mills, RE, Dayama, G, Emery, S, Jun, G, Shi, X, Quitadamo, A, Lunter, G, McVean, GA, Chen, K, Fan, X, Chong, Z, Chen, T,

Witherspoon, D, Xing, J, Eichler, EE, Chaisson, MJ, Hormozdiari, F, Huddleston, J, Malig, M, Nelson, BJ, Sudmant, PH, Parrish, NF, Khurana, E, Hurles, ME, Blackburne, B, Lindsay, SJ, Ning, Z, Walter, K, Zhang, Y, Gerstein, MB, Abyzov, A, Chen, J, Clarke, D, Lam, H, Mu, XJ, Sisu, C, Zhang, J, Zhang, Y, Gerstein, MB, Abyzov, A, Chen, J, Clarke, D, Lam, H, Mu, XJ, Sisu, C, Zhang, J, Zhang, Y, Gibbs, RA, Yu, F, Bainbridge, M, Challis, D, Evani, US, Kovar, C, Lu, J, Muzny, D, Nagaswamy, U, Reid, JG, Sabo, A, Yu, J, Guo, X, Li, W, Li, Y, Wu, R, Marth, GT, Garrison, EP, Leong, WF, Ward, AN, Del Angel, G, DePristo, MA, Gabriel, SB, Gupta, N, Hartl, C, Poplin, RE, Clark, AG, Rodriguez-Flores, JL, Flicek, P, Clarke, L, Smith, RE, Zheng-Bradley, X, MacArthur, DG, Mardis, ER, Fulton, R, Koboldt, DC, Gravel, S, Bustamante, CD, Craig, DW, Christoforides, A, Homer, N, Izatt, T, Sherry, ST, Xiao, C, Dermitzakis, ET, Abecasis, GR, Min Kang, H, McVean, GA, Gerstein, MB, Balasubramanian, S, Habegger, L, Gerstein, MB, Balasubramanian, S, Habegger, L, Yu, H, Flicek, P, Clarke, L, Cunningham, F, Dunham, I, Zerbino, D, Zheng-Bradley, X, Lage, K, Jespersen, JB, Horn, H, Montgomery, SB, DeGorter, MK, Khurana, E, Tyler-Smith, C, Chen, Y, Colonna, V, Xue, Y, Gerstein, MB, Balasubramanian, S, Fu, Y, Kim, D, Gerstein, MB, Balasubramanian, S, Fu, Y, Kim, D, Auton, A, Marcketta, A, Desalle, R, Narechania, A, Sayres, MA, Garrison, EP, Handsaker, RE, Kashin, S, McCarroll, SA, Rodriguez-Flores, JL, Flicek, P, Clarke, L, Zheng-Bradley, X, Erlich, Y, Gymrek, M, Willems, TF, Bustamante, CD, Mendez, FL, Poznik, GD, Underhill, PA, Lee, C, Cerveira, E, Malhotra, A, Romanovitch, M, Zhang, C, Abecasis, GR, Coin, L, Shao, H, Mittelman, D, Tyler-Smith, C, Ayub, Q, Banerjee, R, Cerezo, M, Chen, Y, Fitzgerald, TW, Louzada, S, Massaia, A, McCarthy, S, Ritchie, GR, Xue, Y, Yang, F, Tyler-Smith, C, Ayub, Q, Banerjee, R, Cerezo, M, Chen, Y, Fitzgerald, TW, Louzada, S, Massaia, A, McCarthy, S, Ritchie, GR, Xue, Y, Yang, F, Gibbs, RA, Kovar, C, Kalra, D, Hale, W, Muzny, D, Reid, JG, Wang, J, Dan, X, Guo, X, Li, G, Li, Y, Ye, C, Zheng, X, Altshuler, DM, Flicek, P, Clarke, L, Zheng-Bradley, X, Bentley, DR, Cox, A, Humphray, S, Kahn, S, Sudbrak, R, Albrecht, MW, Lienhard, M, Larson, D, Craig, DW, Izatt, T, Kurdoglu, AA, Sherry, ST, Xiao, C, Haussler, D, Abecasis, GR, McVean, GA, Durbin, RM, Balasubramaniam, S, Keane, TM, McCarthy, S, Stalker, J, Durbin, RM, Balasubramaniam, S, Keane, TM, McCarthy, S, Stalker, J, Chakravarti, A, Knoppers, BM, Abecasis, GR, Barnes, KC, Beiswanger, C, Burchard, EG, Bustamante, CD, Cai, H, Cao, H, Durbin, RM, Gerry, NP, Gharani, N, Gibbs, RA, Gignoux, CR, Gravel, S, Henn, B, Jones, D, Jorde, L, Kaye, JS, Keinan, A, Kent, A, Kerasidou, A, Li, Y, Mathias, R, McVean, GA, Moreno-Estrada, A, Ossorio, PN, Parker, M, Resch, AM, Rotimi, CN, Royal, CD, Sandoval, K, Su, Y, Sudbrak, R, Tian, Z, Tishkoff, S, Toji, LH, Tyler-Smith, C, Via, M, Wang, Y, Yang, H, Yang, L, Zhu, J, Bodmer, W, Bedoya, G, Ruiz-Linares, A, Cai, Z, Gao, Y, Chu, J, Peltonen, L, Garcia-Montero, A, Orfao, A, Dutil, J, Martinez-Cruzado, JC, Oleksyk, TK, Barnes, KC, Mathias, RA, Hennis, A, Watson, H, McKenzie, C, Qadri, F, LaRocque, R, Sabeti, PC, Zhu, J, Deng, X, Sabeti, PC, Asogun, D, Folarin, O, Happi, C, Omoniwa, O, Stremlau, M, Tariyal, R, Jallow, M, Joof, FS, Corrah, T, Rockett, K, Kwiatkowski, D, Kooner, J, HiÃªn, TT, Dunstan, SJ, Hang, NT, Fonnie, R, Garry, R, Kanneh, L, Moses, L, Sabeti, PC, Schieffelin, J, Grant, DS, Gallo, C, Poletti, G, Saleheen, D, Rasheed, A, Saleheen, D, Rasheed, A, Brooks, LD, Felsenfeld, AL, McEwen, JE, Vaydylevich, Y, Green, ED, Duncanson, A, Dunn, M, Schloss, JA, Wang, J, Yang, H, Auton, A, Brooks, LD, Durbin, RM, Garrison, EP, Min Kang, H, Korbel, JO, Marchini, JL, McCarthy, S, McVean, GA, Abecasis, GR, Auton, A, Brooks, LD, Durbin, RM, Garrison, EP, Min Kang, H, Korbel, JO, Marchini, JL, McCarthy, S, McVean, GA, Abecasis, GR **(2015)**. A global reference for human genetic variation. *Nature*, 526, 7571:68-74.
